# High-Resolution Ultrasound and Speckle Tracking: a non-invasive approach to assess *in vivo* gastrointestinal motility during development

**DOI:** 10.1101/2022.02.01.478689

**Authors:** Pierre Sicard, Amandine Falco, Sandrine Faure, Jérome Thireau, Stéphanie E. Lindsey, Norbert Chauvet, Pascal de Santa Barbara

## Abstract

Gastrointestinal motor activity has been extensively studied in adults, conversely only few studies have investigated fetal motor skills. When the gastrointestinal tract starts to contract during the embryonic period and how this function evolves during development are not known. Here, we adapted a non-invasive high-resolution echography technique combined with speckle tracking analysis to examine the gastrointestinal tract motor activity dynamics during chick embryo development. We provided the first recordings of fetal gastrointestinal motility in living embryos. We found that although gastrointestinal contractions appear very early during development, they become synchronized only at the end of the fetal period. To validate this approach, we used various pharmacological inhibitors and *BAPX1* gene overexpression *in vivo*. We found that the enteric nervous system determinates the onset of the synchronized contractions in the stomach. Moreover, alteration of smooth muscle fiber organization led to an impairment of this functional activity. Altogether, our findings show that non-invasive high-resolution echography and speckle tracking analysis allow visualizing and quantifying gastrointestinal motility during development and highlight the progressive acquisition of functional and coordinated gastrointestinal motility before birth.

## INTRODUCTION

In vertebrates, the gastrointestinal (GI) tract is essential for the absorption of water and nutrients. Early during embryo development, the GI tract is formed as a closed primitive and uniform tube, composed of endoderm and mesenchyme, that becomes regionalized along the anterior–posterior (AP) axis into various organs (esophagus, stomach, duodenum, and intestines) (de Santa Barbara et al., 2002; de Santa Barbara et al., 2003). The mesenchyme gives rise (from the outer to the inner part of the gut wall) to the longitudinal and circular muscle layers, the submucosa, and the muscularis mucosae, close to the epithelial lining (Le Guen et al., 2015). The circular and longitudinal smooth muscle layers align in orthogonal orientations to ensure the gut coordinated contraction and relaxation (Huycke et al., 2019; Roberts, 2000). Concomitantly with these morphological events, the GI mesenchyme is colonized by neural crest-derived cells, a cell population that gives rise to the enteric nervous system (ENS), the intrinsic innervation of the GI tract (Burns et al., 2009). The ENS originates predominantly from vagal enteric neural crest-derived cells (vENCDCs) that delaminate from the neural tube, enter the esophageal mesenchyme, and populate the entire GI tract, from the esophagus to the terminal colon, through an AP migration wave (Burns and Douarin, 1998; Burns et al., 2000; Fairman et al., 1995; Faure et al., 2015; Le Douarin and Teillet, 1973; Yntema and Hammond, 1954). During their migration along the GI tract, vENCDCs proliferate and differentiate into the ENS neurons and glial cells. They form two concentric plexuses of ganglion cells; the myenteric plexuses are localized in the GI wall muscle layers and control smooth muscle contraction and relaxation (Bourret et al., 2017; Chevalier, 2018; Heanue et al., 2016).

Vertebrate GI motor activity has been extensively studied in adults, but rarely during embryo development. During the prenatal period, the human GI tract digests and absorbs nutrients from the amniotic fluid and propels the meconium (Mclain, 1963). There are good clinical evidences that in late gestation, fetal growth requires an intact and functional GI tract for swallowing the amniotic fluid and for the enteral uptake of nutrients (Koppen et al., 2017; Singendonk et al., 2014). However, it is not known when and how digestive motor skills appear and develop during development, mainly due to the limitations of *in vivo* embryo assessment. Various invasive approaches using different dissected GI segments for organ culture showed that embryonic GI segments can contract autonomously or upon stimulation (Roberts et al., 2010). These studies demonstrated that the first contractile waves are due to spontaneous smooth muscle contractions (Chevalier, 2018; Roberts et al., 2010). The transition from uncoordinated to rhythmic motility patterns in the developing intestine has been associated with the activity of ENS (Chevalier et al., 2019; Roberts et al., 2010) and Interstitial Cells of Cajal (ICCs) (Chevalier et al., 2020), which are mesenchymal cells that have differentiated from digestive mesenchymal progenitors common to smooth muscle cells and ICCs (Guérin et al., 2020; Le Guen et al., 2015).

Therefore, a non-invasive *in vivo* approach that can reproducibly identify, quantify, and follow GI motility during fetal development is needed to identify the implicated physiological key mechanism(s) and monitor the emergence of motility. This will allow scientists to understand how rhythmic motility patterns are set up and their alterations in infants with functional GI disorders (Martire et al., 2021; Thapar et al., 2018). Here, we used high-frequency ultrasound imaging and speckle tracking analysis to study GI motility in chick embryos. This method allowed us to identify *in vivo* early GI motility, its changes and functional profiles during chick embryo development. This approach combined with pharmacological inhibitors of smooth muscle contraction and of ENS or ICC activity allowed us to highlight ENS role in the synchronization of stomach contractions *in vivo*. Moreover, by overexpressing *BAPX1* in the stomach mesenchyme *in vivo*, we demonstrated that smooth muscle fiber organization is essential for functional fetal GI motility.

## RESULTS

### Stomach contraction undergoes dynamic changes to reach coordinated patterns during chick embryo development

To enable the non-invasive *in vivo* investigation of the GI tract in chick embryos, we used the high-resolution echography imaging technique that was previously applied to monitor heart changes during chick embryo development (McQuinn et al., 2007). At embryonic day 15 (E15), the latest stage we analyzed, we could visualize all embryonic GI segments (stomach, small intestine, and colon) and their associated organs (liver, lung, pancreas). Besides the anatomical structures, we also monitored the stomach deformations associated with the dynamic opening and closure of its lumen (Fig. 1A, Movie 1). We observed these movements in small intestine and colon (Movies 2&3) as well. We then used high-resolution echography to investigate the onset and changes of stomach motility during embryogenesis, from E8 to E15. We could visualize the stomach structure at E8, and unexpectedly, we observed stomach movements already at this stage (Fig. 1B, Movie 4). After recording images of the GI for 1 minute at 25 images/sec, we used the speckle tracking analysis software (VevoLab) to analyze them. Strain was defined as the relative change in length, and was determined with the formula ε = (L-L_0_)/L_0_ where L_0_ is the baseline length and L is the length at maximum contraction. Using this analysis, we generated a deformation map (3D strain) at E8 and quantified the stomach deformation (ranking from -9.5 to +10.3). As we observed asynchronous deformations around the stomach circumference, suggesting that contractions began in an uncoordinated manner (Fig. 1C). At E13 (Fig. 1B, Movie 5), the magnitude of stomach deformation increased (from -24.7 to +21.8), but became increasingly confined to specific zones (Fig. 1C). At E15, the stomach deformation percentage was still elevated (from -8.7 to +21.9). Moreover, we detected these deformations confined to specific zones (Fig. 1C), highlighting that stomach contraction evolves during development. Using high-resolution echography, we found that the stomach area increased rapidly (by 13-fold) between E8 and E15 (Figure 1D): from 2.327 mm^2^ at E8 to 21.41 mm^2^ at E13, and 30.47 mm^2^ at E15. To quantify stomach deformation changes between E8 and E15, we used the speckle-tracking strain curves obtained from high-resolution echography images. We quantified stomach strain dyssynchrony based on the maximal radial time-strain curves and time-strain curves of six delineated stomach segments (Fig. 1E), adapting a commonly used cardiac resynchronization index (Suffoletto et al., 2006), to the gastrointestinal system. We found that dyssynchrony was very high at E8 (2092.58), decreased at E13 (930.79), and was near the basal line (230.89) at E15, indicating high movement synchronization at E15. The dyssynchrony values at E13 and E15 were significantly different (Figure 1F; P<0.05). Altogether, the high-resolution echography approach demonstrated *in vivo* the dynamic contraction pattern of stomach that changed to synchronized movements at E15.

**Figure 1:**
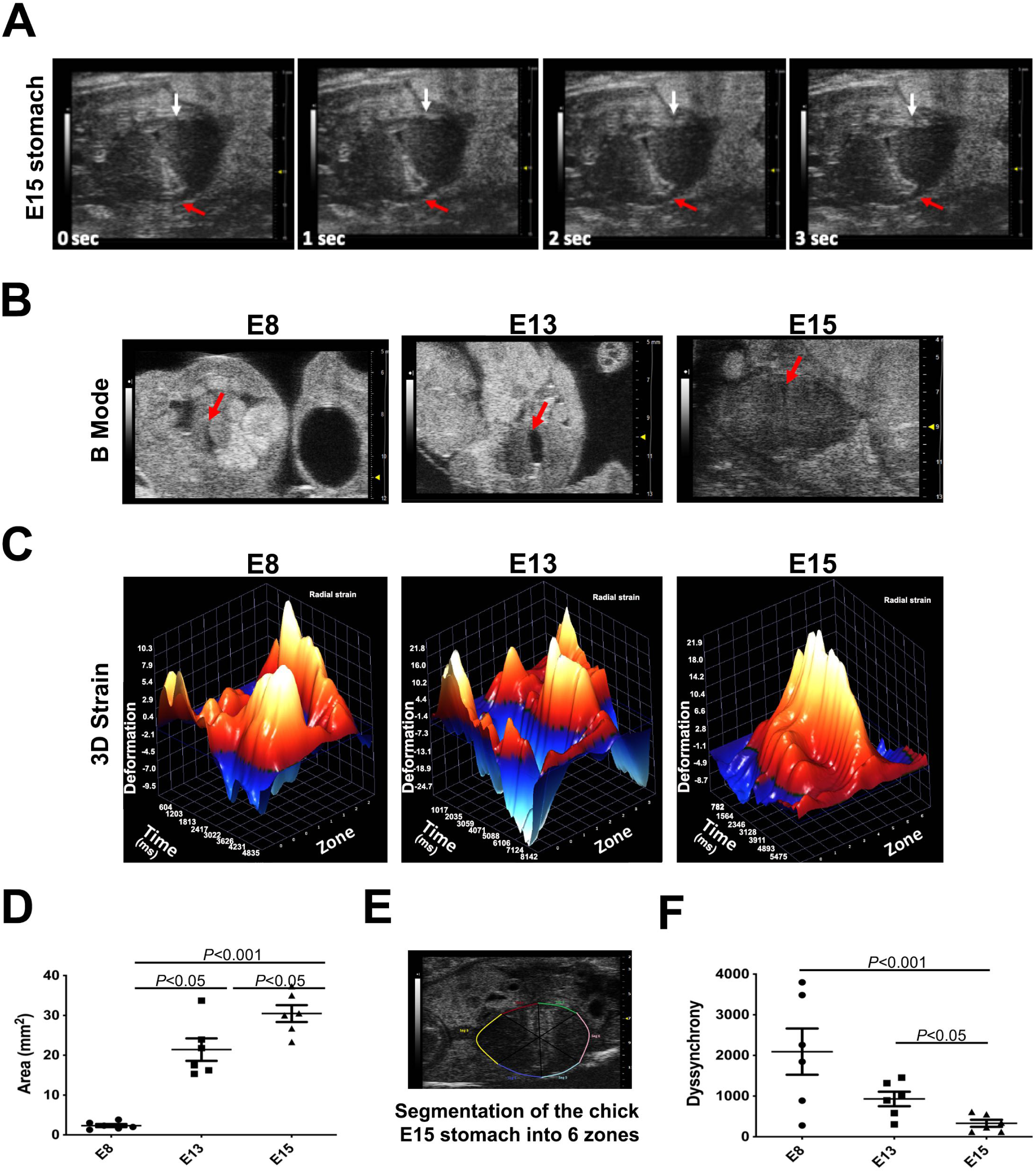
Dynamic contractile activity during fetal chick stomach development. (A) Observation of stomach contractions *in vivo* using high-resolution echography in E15 chick embryos. White arrows, muscle contraction waves; red arrows, lumen movement. (B) Stomach anatomy *in vivo* analysis using high-resolution echography at different stages of chick embryo development (E8, E13, and E15). Red arrows, stomach lumen. (C) 3D strain analysis of the stomach contractile activity *in vivo* using high-resolution echography imaging data during chick embryo development (E8, E13, and E15). (D) Quantification of the stomach area at the indicated developmental stages (n=6/stage). (E) Segmentation of the chick embryo stomach into six zones to monitor deformation. (F) Evaluation of dyssynchrony at the indicated developmental stages (n=6/stage).

### During development, colon and stomach motility coordination patterns appear later than in small intestine

Most previous studies on GI motility onset focused on the small intestine using organs isolated from mouse and chick embryos (Chevalier, 2018; Chevalier et al., 2020; Roberts et al., 2010). Using high-resolution echography, we could monitor the small intestine from E13 onwards, and we detected a rhythmic movement already at this stage (Fig. 2A, B). The small intestine area increased from 0.1825 mm^2^ at E13 to 0.385 mm^2^ at E15 (Fig. 2C; P <0.05). Using speckle tracking analysis, we measured the circumferential and radial small intestine strains (i.e. change in length over the original length) (Fig. 2D). The absence of a significant difference in the percentages of small intestine radial (9.32% at E13 and 11.63% at E15) and circumferential (15.91% at E13 and 15.15% at E15) strains indicated no evolution of the small intestine contraction between E13 and E15 (Fig. 2E). We then observed the colon lumen in the longitudinal orientation, and detected dynamic movements with the presence of rhythmic waves (Fig. 2F; Movie 3). The colon diameter slightly increased from 1.223 mm at E13 to 1.343 mm at E15 (Figure 2H, not significant). Using speckle tracking analysis, we measured the colon longitudinal strain (Fig. 2G), and found that the longitudinal deformation was significantly increased at E15 compared with E13 (1.525% at E13 and 4.025% at E15) (Fig. 2H; P <0.05). Altogether, the high-resolution echography approach showed efficient small intestine motility already at E13, whereas the colon motor skills continued to progress from E13 to E15.

**Figure 2:**
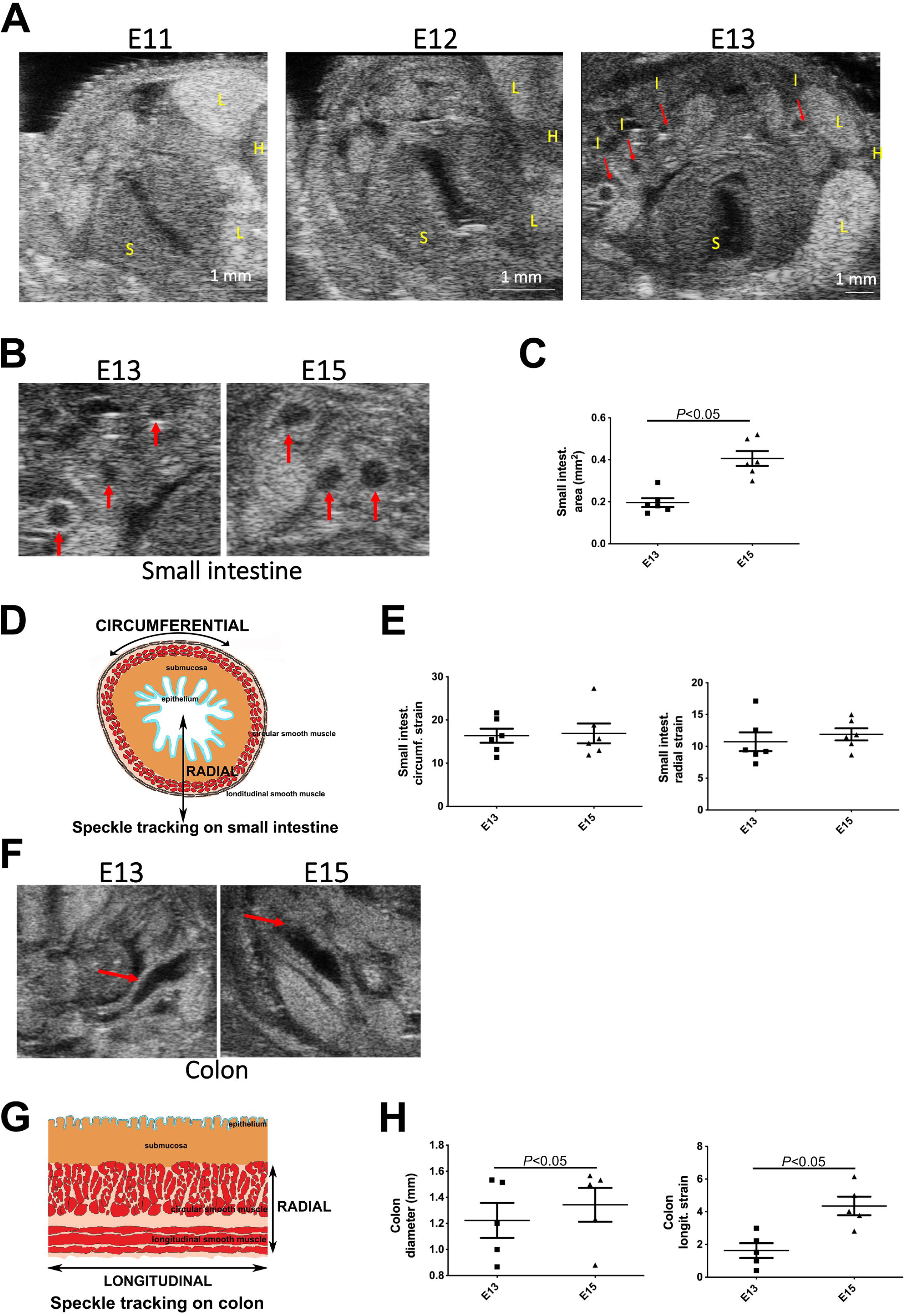
Dynamic contractile activity during chick embryo small intestine and colon development. (A) Small intestine anatomy *in vivo* using high-resolution echography during chick embryo development: E11 (stage 37), E12 (stage 38), and E13 (stage 39). Red arrows, small intestine lumen. H, heart; I, small intestine; L: lung; S, stomach. (B) Small intestine anatomy by high-resolution echography at E13 and E15. Red arrows, small intestine lumen. (C) Quantification of the small intestine area at the indicated developmental stages (n=6/stage). (D) Schematic representation of the speckle tracking analysis of the small intestine during chick embryo development. (D) Quantification of small intestine circumferential and radial strains at the indicated developmental stages (n=6/stage). (F) Colon anatomy analysis by high-resolution echography at E13 and E15. Red arrows, colon lumen. (G) Schematic representation of the speckle tracking analysis of colon during chick embryo development. (H) Quantification of colon diameter and longitudinal strain at the indicated developmental stages (n=6/stage).

### The enteric nervous system is implicated in fetal stomach contractions

To challenge our high-resolution echography approach and to determine the origin of stomach contractions, we developed an approach to deliver in the GI lumen drugs that target specific cell types (smooth muscle cells, enteric neurons, and ICCs) found in the developing stomach musculature (Fig. 3A). Using high-resolution echography, we monitored stomach contractions at E15 before and 1 hour after drug delivery. As a control, we also determined the heart rate. We first used the calcium channel blocker cobalt chloride (CoCl_2_) to inhibit the autonomous smooth muscle contraction. We used speckle tracking analysis, to measure the circumferential and radial stomach strains (i.e. change in length over the original length) (Fig. 3B). Results showed that CoCl_2_ led to a decrease in stomach circumferential velocity (from 6.514 mm/sec before to 2.828 mm/sec after treatment, P <0.05) and radial displacement (from 0.1695 mm before to 0.082 mm after treatment, P <0.05) (Fig. 3C). Moreover, we confirmed that this approach specifically targets the stomach because heart rate was not altered by CoCl_2_ (187.66 beat/min before and 188.8 beat/min after treatment, not significant) (Fig. 3C). These data demonstrated that the measured deformations were specific to the intrinsic stomach smooth muscle contractions. To evaluate ENS contribution to stomach contraction, we used the sodium neural channel blocker tetrodotoxin (TTX). We found that TTX decreased the stomach circumferential velocity (from 7.464 mm/sec before to 4.44 mm/sec after treatment, P <0.05) and radial displacement (from 0.1518 mm before to 0.032 mm after treatment, P <0.05) (Fig. 3D), leading to stomach contraction inhibition. The heart rate remained constant (185.6 beat/min before and 181.5 beat/min after treatment, not significant) (Fig. 3D). We then used Imatinib to block ICC activity (Beckett et al., 2007; Chevalier et al., 2020; Kim et al., 2010; Popescu et al., 2006). After 1 hour, we observed a decrease in stomach circumferential velocity (from 9.566 mm/sec before to 3.0336 mm/sec after treatment, P <0.0001) and radial displacement (from 0.237 mm before to 0.064 mm after treatment, P <0.001) (Fig. S1), leading to stomach contraction inhibition. However, as Imatinib also affected the heart rate, which decreased from 212 beat/min before to 148.4 beat/min after treatment, (P <0.05), results do not reflect a targeted ICC role in stomach contraction regulation. This data shows that high-resolution echography allows for monitoring of GI tract contractions *in vivo*, facilitating robust quantitative analysis. Moreover, using GI-targeted drug delivery, we found that at E15 the ENS contributes to stomach motility regulation.

**Figure 3:**
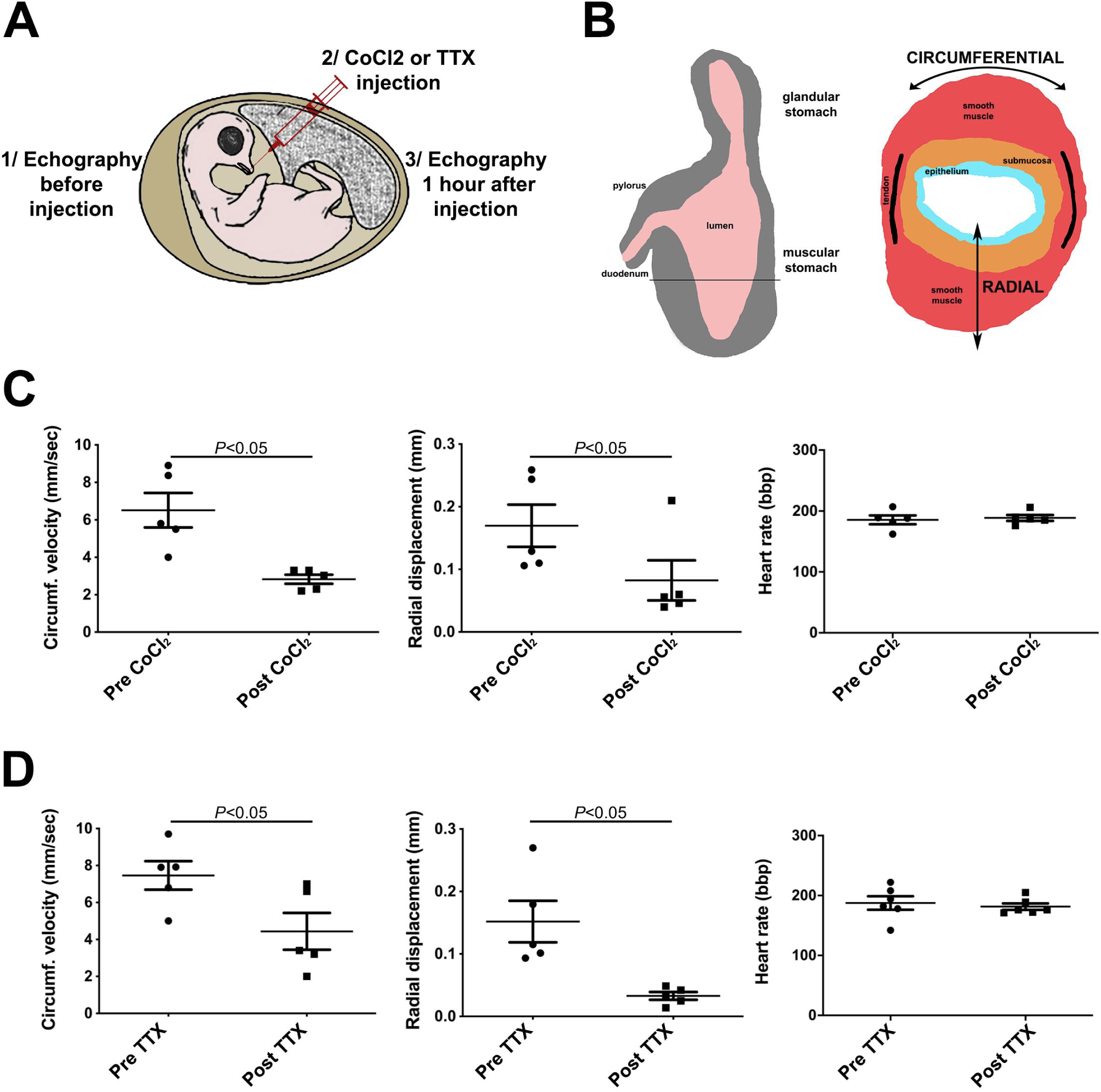
Monitoring stomach contractile activity in E15 chick stomach. (A) Schematic representation of the method used to target GI tract with drugs. (B) Schematic representation of the speckle tracking analysis of stomach during chick embryo development. (C) Effect of cobalt chloride (10 µM; CoCl_2_) on stomach circumferential strain velocity, stomach radial strain changes, and heart rate in E15 chick embryos (n=5). (D) Effect of tetrodotoxin (25 µM; TTX) on stomach circumferential strain velocity, stomach radial strain changes, and heart rate in E15 embryos (n=5).

### Smooth muscle layer organization is essential for fetal stomach contraction coordination

The development of an approach to monitor digestive contractility *in vivo* opens the way to study the role of specific genes. BMP signaling activity has been involved in the development and differentiation of the digestive mesenchyme into smooth muscle (De Santa Barbara et al., 2005; Notarnicola et al., 2012). More recently, it has been shown that BMP activity modulation in the small intestine leads to the alteration of the longitudinal smooth muscle orientation relative to the circular layer (Huycke et al., 2019). Upstream of the BMP ligand, the homeobox gene *BAPX1* (also known as *NKX3*.*2*) is expressed in the vertebrate distal stomach mesenchyme (Faure et al., 2013; Nielsen et al., 2001; Verzi et al., 2009). BAPX1 negatively regulates *BMP4* expression and consequently BMP activity in the stomach mesenchyme, resulting in the expansion of the gastric and duodenal mesenchyme mass (De Santa Barbara et al., 2005; Nielsen et al., 2001). However, *BAPX1* functions in the stomach smooth muscle differentiation have not been investigated yet. Therefore, to evaluate the impact of BMP activity modulation on fetal stomach contractions, we used the avian replication-competent retroviral misexpression system to continuously express *BAPX1* in the developing stomach mesenchyme (Faure et al., 2015; McKey et al., 2016). Then, high-resolution echography showed that the *BAPX1*-expressing stomach lumen of E13 chick embryos was devoid of refringent content compared with E13 *GFP-*expressing stomach lumens (controls) (Fig. 4A). Moreover, *BAPX1*-expressing stomachs were hypotonic compared with controls and did not show a rhythmic muscular deformation and displacement (Movies 6 and 7). Using speckle tracking analysis, we found that the stomach circumferential velocity (8.202 mm/sec for *BAPX1-*expressing and 5.734 mm/sec for control stomachs, P <0.05) and radial displacement (0.1977 mm for *BAPX1-*expressing and 0.082 mm for control stomachs, P <0.01) (Figure 4B) were decreased in E13 *BAPX1*-expressing stomachs, leading to stomach contraction inhibition. We then evaluated *BAPX1* expression effect on the stomach morphology and smooth muscle differentiation status. *BAPX1* overexpression induced minor morphological defects in the distal and proximal stomach (respectively gizzard and proventriculus in birds) at E13 (Fig. 4C, upper panels). We confirmed the mesenchyme targeting by *BAPX1*- and *GFP*-expressing (control) retroviruses with anti-gag antibodies (Fig. 4C, lower panels). We detected the smooth muscle marker gamma smooth muscle actin (γSMA) by immunostaining in transversal paraffin sections of E13 control- and *BAPX1*-expressing stomachs, suggesting that the induction of contractile proteins in smooth muscle was not impaired (Fig. 4D). However, in *BAPX1*-expressing stomachs, the fiber structure was disorganized (compare white and red arrows). Expression of the pan-neuronal (B3-TUBULIN (TUJ1)) marker indicated that enteric neurons were present in both conditions (Fig. 4D). To better characterize the smooth muscle organization, we modified and optimized the RapidClear® tissue-clearing protocol to obtain whole translucent stomachs. We assessed the 3D impact of *BAPX1* overexpression using light-sheet microscopy. In E13 control stomach samples, 3D analysis of γSMA expression highlighted the presence of several muscle bundles organized in parallel fibers with an orthogonal orientation in the most posterior part of the stomach (Fig. 4E, white arrow in the dorsal view; Movie 8). Like in control samples, in E13 *BAPX1*-expressing stomach samples, muscle bundles were organized in parallel fibers. However, the orthogonal orientation of the second muscle layer was altered: fibers were present, but harbored multiple orientations (Fig. 4E, red arrow in the ventral view; Movie 9). *BAPX1* expression also led to the inversion of the muscle bundle orientation (Fig. 4E, compare red and white arrows in the dorsal view). The changes in the muscle fiber orientation observed in whole *BAPX1*-expressing stomach samples were confirmed in virtual sections (Fig. 4F, compare red and white arrows). As the intestinal smooth muscle differentiation contributes to the ENS organization (Graham et al., 2017), we examined also the neuronal network. The ENS network spread in the smooth muscle layer sparing the tendons (Fig. 4E,F; Movie 8) (Le Guen et al., 2009). However, the neuronal mesh was less dense and less interconnected in E13 *BAPX1*-expressing stomachs where we observed disorganized smooth muscle bundles (Fig. 4E,F; Movie 9). Altogether, the combination of high-resolution echography *in vivo* imaging, gain of function approach, and 3D tissue-clearing immunofluorescence allowed us to demonstrate that BMP signaling deregulation in the stomach mesenchyme alters the segmental orientation of the smooth muscle layers, a feature associated with impaired fetal motility.

**Figure 4:**
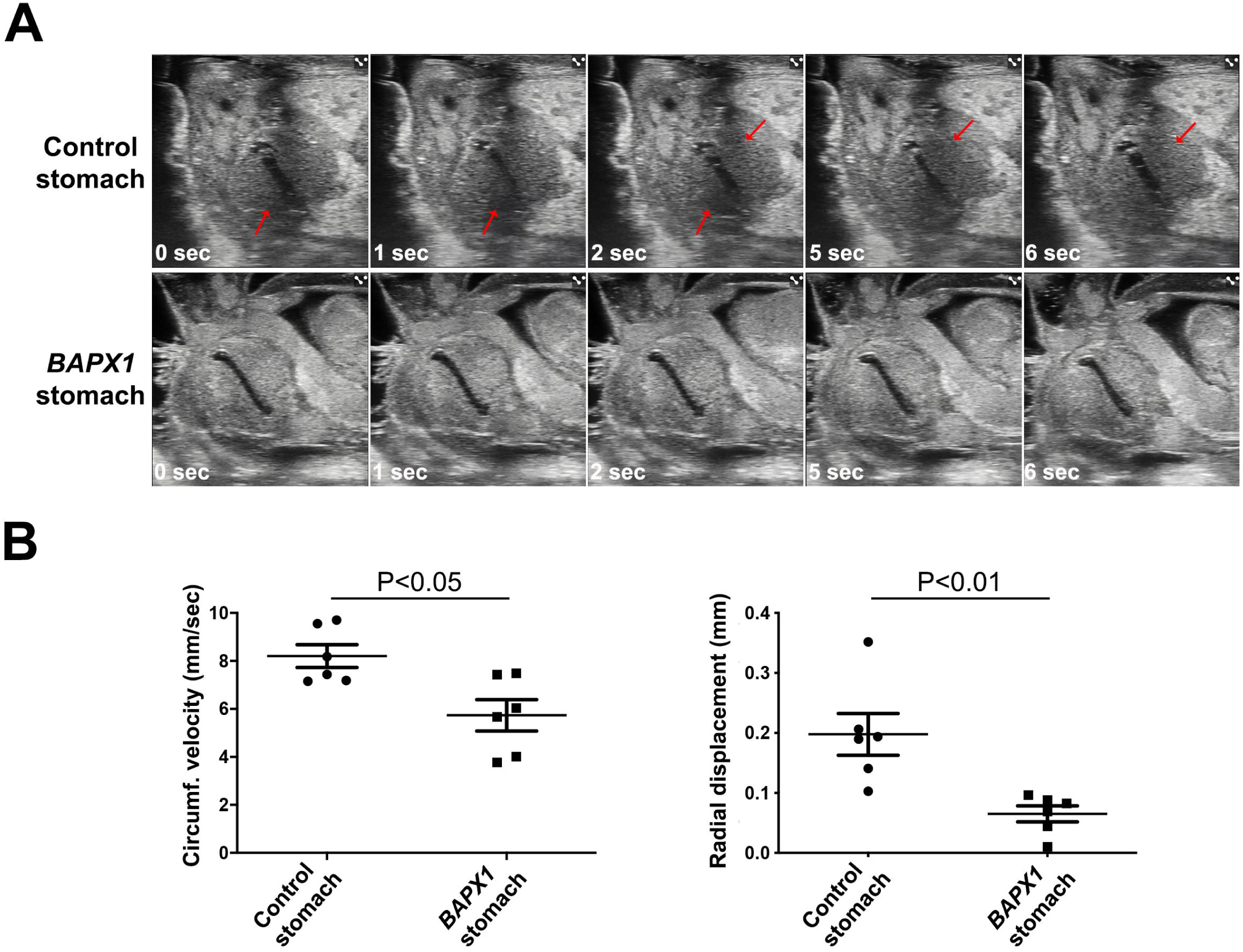
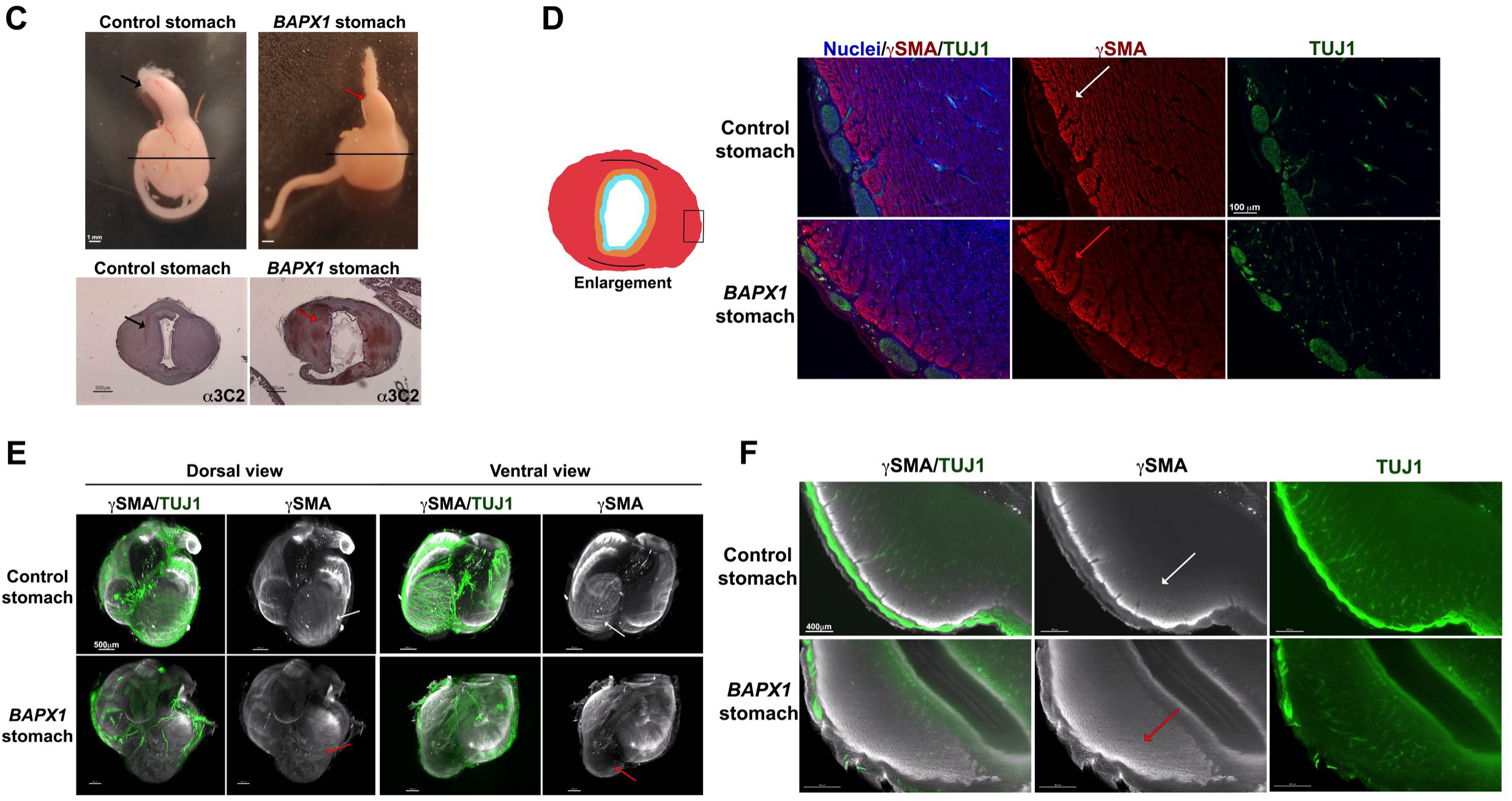
Interfering *in vivo* with the BMP signaling pathway affects stomach contractility in chick embryos. (A) Observation of stomach contractions *in vivo* using high-resolution echography in E13 Control and *BAPX1-*overexpressing stomachs. Red arrows, muscle movement in the E13 control stomach. (B) Speckle tracking analysis to evaluate the impact of BAPX1 expression on stomach circumferential strain velocity and stomach radial strain changes in E13 chick embryos (n=6). (C) Effect of BAPX1 expression on E13 *BAPX1*-expressing stomach gross morphology compared with E13 Control stomach (upper panels; ventral view). Red and black arrows indicate the resulting proventriculus. Transversal paraffin sections of E13 Control and *BAPX1*-expressing stomachs analyzed by immunohistochemistry with an anti-gag (3C2) antibody (lower panel). Red arrows indicate *BAPX1* expression in the E13 *BAPX1*-expressing stomach. Scale bars: 500µm. (D) Transversal paraffin sections of E13 *BAPX1*-expressing and Control stomachs close to the epithelium (left panel). Nuclei were visualized with Hoechst. Antibodies against smooth muscle cells (γSMA), and neuronal cells (TUJ1) were used. Scale bars, 100 µm. (E) Light-sheet microscopy analysis after RapidClear tissue clearing and γSMA and TUJ1 immunofluorescence staining of E13 whole Control and *BAPX1*-expressing stomachs. Scale bars: 500 µm. White and red arrows indicate smooth muscle fiber organization in Control and *BAPX1*-expressing stomachs. (F) Virtual longitudinal sections of whole E13 *BAPX1*-expressing and Control stomachs stained for γSMA and TUJ1. White and red arrows indicate difference in circular muscle fiber orientation. Scale bars, 400 µm (not in my figure).

## Discussion

Using an approach that combines *in vivo* non-invasive high-resolution echography imaging and speckle-tracking analysis, we characterized GI motility patterns during chick embryo development. We adapted and validated this approach that is mainly used in cardiovascular physiopathology for perinatal gastroenterology investigations using chick embryos, a vertebrate model that allows the longitudinal investigation of all GI domains. Until now, embryonic gut motility has been studied only in organ culture systems. However, depending on the GI segment and preparation method (open, tubular, muscle strips), the motility patterns and contractile behavior can be biased by the apparatus used to measure smooth muscle contractility (Barnes et al., 2014). Our approach demonstrated though an in-vivo approach that the early erratic contractions of the stomach musculature observed at E8 became synchronized at E15. Investigation of other segments of the developing GI tract highlighted differences in motility onset along the AP axis. Like the stomach, the colon motility became fully efficient at E15. Conversely, the small intestine motility was effective at E13, as indicated by the presence of efficient waves of peristalsis. The timing of efficient intestinal motility coincides with the appearance of the longitudinal smooth muscle layer in the chick small intestine (Shyer et al., 2013), suggesting the requirement of the second smooth muscle layer for efficient contractions. The onset of GI smooth muscle differentiation and its regional differences have been previously described in the chick embryo (Bourret et al., 2017; Graham et al., 2017). Conversely, the precise timing of the sequential differentiation of the distinct smooth muscle layers along the AP of the GI tract was unknown. Our functional observations support regional differences in the appearance of the longitudinal smooth muscle layer.

Stomach development and its specific morphogenesis have been extensively studied in several animal models (for review, (Grapin-Botton, 2005; Kim and Shivdasani, 2016; Le Guen et al., 2015; McLin et al., 2009)), but few studies have addressed the development of motor skills in organ culture, and even fewer *in vivo*. The human stomach musculature consists of two smooth muscle layers, an external longitudinal and internal circular muscle layer, for most of its extent. In addition, there is an oblique muscle layer, internal to the circular muscle layer, in the gastro-esophageal junction region (Di Natale et al., 2021; Hur, 2020). Adult gastric movements depend on the generation of electrical rhythmicity and electrical conduction. The directions and strengths of the forces generated when the muscle is excited, depend on the musculature organization. However, there is no detailed quantitative data on the vertebrate stomach musculature organization, although their innervation has been extensively described (Furness et al., 2020). Using the high-resolution echography/speckle tracking approach and pharmacological inhibitors, we found that stomach contractions are controlled by enteric neurons at E15. This suggests the importance of the establishment of stomach contraction to ensure the fetal transit that is also essential for the fetus growth. Using a genetic approach, we then showed that deregulation of the BMP pathway activity affects the organization of the longitudinal smooth muscle layer and its orientation relative to the circular layer, but not gastric smooth muscle differentiation and the oblique smooth muscle layer. The high-resolution echography/speckle tracking approach in *BAPX1*-expressing embryos allowed us to detect *in vivo* a functional alteration that impaired contraction, despite the effective smooth muscle cell differentiation, highlighting the importance of this new approach.

Our data demonstrated differences in peristalsis onset along the AP axis at a developmental stage that in the human embryo corresponds to 12 and 14 weeks of gestation. The pediatric Chronic Intestinal Pseudo-Obstruction (CIPO) syndrome is the most severe functional gastrointestinal disorder (high morbidity and mortality rates), but lacks standardized diagnostic and therapeutic approaches (Thapar et al., 2018). Recently, it was reported that the *ACTG2* gene, which encodes γSMA, is mutated in 30% of children with CIPO who have a worse outcomes and severe GI dysmotility often associated with the presence of megacystis (Hashmi et al., 2021; Matera et al., 2016). In pediatric patients with CIPO, prenatal signs are detected only in about 20% of cases, mainly the presence of megacystis, although 50–70% of infants show clinical signs in the first month after birth (Di Nardo et al., 2017). Multi-visceral dilation has been observed, using standard ultrasonography, in two fetuses with CIPO (Shen et al., 2007). This suggests that the non-invasive high-resolution echography approach could be useful to easily and routinely assess digestive function in fetuses before birth. However, routine ultrasound examination to detect GI dysmotility is not part of the recent antenatal recommendations (Thapar et al., 2018).

Altogether, we demonstrated that GI contractions occur during fetal development and progress from an uncoordinated pattern to a more powerful coordinated profile, in agreement with the hypothesis that mechanical features of smooth muscle and/or their environment, such as intrinsic and extrinsic innervation, influence embryonic peristalsis.

## MATERIALS AND METHODS

### Animal model and ultrasound data acquisition and analysis

Fertilized chicken eggs (Les Bruyères, Dangers, France) were incubated at 38^°^C in a humidified incubator (SMA Coudelou, France) until use. Although experiments using chick embryos do not require approval by an ethic committee (European law, article 2016/63/UE), they were performed in accordance with the INSERM and CNRS ethics guidelines for animal experimentation. After 2 days of incubation, 4 ml of albumin was removed and a hole was made and sealed with tape to avoid the formation of vessels on the shell top. Briefly, eggs were taken out of the incubator and held under a bright light to localize the embryo. Then, a window of approximately 1.5 by 1.5 cm was sawn in the eggshell. The eggshell and outer membrane were removed to visualize the embryo that was staged by microscopic examination according to Hamburger and Hamilton (Hamburger and Hamilton, 1951) and Southwell (Southwell, 2006). Embryos that were dysmorphic or showed visible bleeding were excluded. E8 (stage 34), E13 (stage 39), and E15 (stage 41) embryos were studied *in vivo* (n=6 embryos per stage). At these stages, embryos float on the left side in the egg yolk. During ultrasonography, each egg was positioned in a dry block heater filled with sand to maintain the temperature of 38°C inside the egg. Vevo2100 and Vevo3100 (Visualsonics) ultrasound system with 40 MHz probes (MS550D and MX550D) were used for *in vivo* image acquisition (spatial resolution of 40 µm). Briefly, the probe was delicately positioned on the eggshell window using an adjustable stand and transducer mount. B-Mode was used to record 2D images of the GI tract for 1 minute at 25 images/sec. The acquisition was repeated three times. B-mode videos were transferred to ImageJ to measure the cross-sectional area and diameter. The speckle tracking analysis software (VevoLab 5.6.1) was used off-line to quantify the maximum regional (stomach, intestine and colon) velocity, displacement, and strain from E8 to E15. Strain is defined as the relative change in length, and is determined with the formula ε = (L-L_0_)/L_0_ where L_0_ is the baseline length and L is the length at maximum contraction. Circumferential strain identifies the contraction along the circular outline, radial strain the thickening of the organ, and longitudinal strain the change in the length relative to the original organ length. Statistical analyses were done with the Prism 8 software and one-way ANOVA with Tukey’s multiple comparison test and a single pool of variance.

### Pharmacological inhibition

To delivery specific compounds in the GI tract, an intra-oral administration technique was developed. To validate this approach, Evan Blue solution was deposited directly in the E15 chick embryo beak with a capillary pipette and 1 hour later the dye was detected in the stomach lumen (data not shown). The sodium channel blocker TTX (1 mM stock solution, Tocris, Bristol, UK) was used to block neurotransmission (D’Antona et al., 2001), and the calcium channel blocker CoCl_2_ (0.1 M stock solution, Sigma-Aldrich, France) was used to block smooth muscle contraction (de Moraes and Carvalho, 1969). Imatinib mesylate (STI571) (10 mM stock solution, Euromedex, France) was used to block KIT and ICC activity (Beckett et al., 2007; Chevalier et al., 2020). Drugs were diluted to the final concentration by adding sterile PBS. Before drug administration, the stomach and heart of each E15 embryo were evaluated by high-resolution echography. This was followed by intra-oral administration of 100 µl of 25 µM TTX, 100 µl of 10 µM cobalt chloride, 100 µl of 20 µM imatinib, or 100 µl of PBS (control). Using fine tools and syringes, the solution was dropped in the beak, and each egg was put back in the incubator to allow the drug progression to the GI tract. After 1 hour, each egg was evaluated by high-resolution echography, as previously described. The unpaired *t-*test was calculated with the Prism 8 software.

### Avian retroviral misexpression system, RapidClear® tissue clearing, and light-sheet microscopy analysis

Fertilized White Leghorn eggs from Morizeau Farm (France) were incubated at 38°C in humidified incubators. The vector to produce replication-competent retroviruses (RCAS) was previously described (Le Guen et al., 2009; Moniot et al., 2004). The DF-1 chicken fibroblast cell line (ATCC-LGC) was transfected with RCAS-based constructs to produce retroviruses that express GFP (Moniot et al., 2004) or BAPX1 (De Santa Barbara et al., 2005; Nielsen et al., 2001). *GFP*-expressing retroviruses alone (as control) or a mix of *BAPX1*+*GFP*-expressing retroviruses were injected in the splanchnopleural mesoderm of stage 10 chick embryos to target the stomach mesenchyme (Le Guen et al., 2009; Moniot et al., 2004; Notarnicola et al., 2012; Roberts et al., 1998). Eggs were then placed at 38°C until high-resolution echography followed by dissection. Only GFP-positive stomachs were analyzed by high-resolution echography and underwent RapidClear® tissue clearing.

For immunofluorescence of paraffin sections, stomachs were gradually dehydrated in ethanol and embedded in paraffin; 10-µm sections were cut using a microtome and collected on poly-L-lysine-coated slides (Thermo Fisher) for immunofluorescence (Faure et al., 2013) using rabbit anti-γSMA (MyBioSource, MBS820899, 1:500 dilution), and mouse anti-TUJ1 (Covance, 1:800 dilution) antibodies. Nuclei were stained with Hoechst 33342 (Molecular Probes). Cells were imaged with a Zeiss AxioVision fluorescence microscope using standard filters, or a ZEISS LSM800 confocal laser-scanning microscope.

For tissue clearing, GI tissues were fixed at room temperature (RT) on an orbital shaker in 4% paraformaldehyde for 2 hours, and then washed in PBS for 1 hour. Samples were transferred to 2% Triton X-100 in PBS solution (containing 0.05% sodium azide) for permeabilization at RT on orbital shaker for 1-2 days, and the washed 3 times for 15min in PBS at RT. Samples were incubated at 4°C on an orbital shaker in blocking solution (10% normal donkey serum, 1% Triton-X100, 0.2% sodium azide in PBS) for 2 days, followed by incubation with primary antibodies (rabbit anti-γSMA (1:300 dilution), and mouse anti-TUJ1 (Covance, 1:200 dilution) in antibody dilution buffer (1% normal donkey serum, 0.2% Triton-X100, 0.2% sodium azide in PBS) at 4°C on orbital shaker for 3-4 days. Samples were washed on orbital shaker 3 times at RT for 1 hour, and at 4°C with washing buffer (3% NaCl, 0.2% Triton-X100 in PBS) overnight. Samples were incubated with anti-rabbit Alexa Fluor 647 (A31573 Invitrogen, 1:300 dilution) and anti-mouse Alexa Fluor 568 (A10037 Invitrogen, 1:300 dilution) secondary antibodies, in dilution buffer at 4°C on an orbital shaker for 2 days. Samples were washed on an orbital shaker at RT 3 times for 1 hour and at 4°C with washing buffer overnight. This was followed by three washes with PBS for 15 minutes/each and sample clearing with RapidClear® at RT overnight. Cleared specimens were placed into 2,2’-Thiodiethanol solution (#166782 Sigma) and tissues were imaged using light-sheet microscopy (UltraMicroscope Blaze, Miltenyi Lavision BioTec). Images were analyzed with Imaris.

## Abbreviations used in this paper

AP: anterior–posterior
CIPO: chronic intestinal pseudo-obstruction
CoCl_2_: cobalt chloride
E: embryonic day
ENS: enteric nervous system
γSMA: gamma smooth muscle actin
GI: gastrointestinal
ICC: interstitial cell of Cajal
RCAS: replication-competent retroviruses
TTX: tetrodotoxin
vENCDC: vagal enteric neural crest-derived cell

## Acknowledgments

The authors thank the members of the “Development of visceral smooth muscle and associated pathologies” team (PhyMedExp), Patrice Bideaux for technical assistance, Emilie Josse for graphical art, the Imagerie du Petit Animal de Montpellier (IPAM, Biocampus Montpellier) for access to high-resolution ultrasound (LRQA Iso9001; France Life Imaging (grant ANR-11-INBS-0006); IBISA; Fondation Leducq (Grant RETP), I-Site Muse) and Montpellier Ressources Imagerie for access to light-sheet microscopy (M-P. Blanchard, A. Sarrazin and O. Faklaris, MRI, Biocampus, Montpellier) facilities.

## SUPPLEMENTAL FIGURES

**Supplementary Figure S1.**
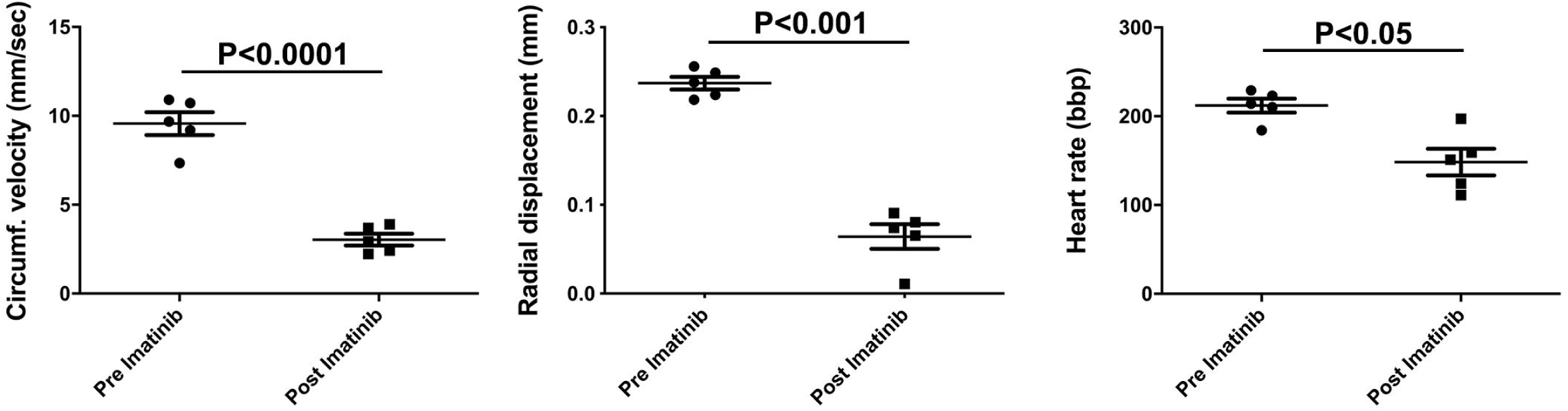
Effect of imatinib. (20 µM) on stomach circumferential strain velocity, stomach radial strain changes, and heart rate in E15 chick embryos (n=5).

## MOVIES

**Movie 1:** Movie obtained using the Vevo2100 ultrasound system to visualize stomach motility at E15.

**Movie 2:** Movie obtained using the Vevo2100 ultrasound system to visualize small intestine motility at E15.

**Movie 3:** Movie obtained using the Vevo2100 ultrasound system to visualize colon motility at E15.

**Movie 4:** Movie obtained using the Vevo2100 ultrasound system to visualize stomach motility at E8.

**Movie 5:** Movie obtained using the Vevo2100 ultrasound system to visualize stomach motility at E13.

**Movie6:** Movie obtained using the Vevo2100 ultrasound system to visualize stomach motility of E13 Control-stomach.

**Movie 7:** Movie obtained using the Vevo2100 ultrasound system to visualize stomach motility of E13 *BAPX1*-overexpressing stomach.

**Movie 8:** 3D Control stomach smooth muscle and ENS imaging at E13 using γSMA and TUJ1 immunofluorescence analysis.

**Movie 9:** 3D *BAPX1*-overexpressing stomach smooth muscle and ENS imaging at E13 using γSMA and TUJ1 immunofluorescence analysis.

